# Whole-gut spatial genomic analysis reveals molecular regionalization of the differentiating zebrafish enteric nervous system

**DOI:** 10.1101/2025.04.17.649413

**Authors:** Rodrigo Moreno-Campos, Nikhita S. Mummaneni, Rosa A. Uribe

## Abstract

The enteric nervous system (ENS) is the intrinsic nervous system of the gut and controls essential functions, such as gut motility, intestinal barrier function, and water balance. The ENS displays a complex 3D architecture within the context of the gut and specific transcriptional states needed to control gut homeostasis. During development, the ENS develops from enteric neural progenitor cells (ENPs) that migrate into the gut and differentiate into functionally diverse neuron types. Incorrect ENS development can disrupt ENS function and induce various gut disorders, including the congenital disease Hirschsprung disease, or various other functional gut neurological disorders, such as esophageal achalasia. In this study, we used the zebrafish larval model and performed whole gut spatial genomic analysis (SGA) of the differentiating ENS at cellular resolution. To that end, a pipeline was developed that integrated early and late developmental ENS stages by linking various spatial and transcriptional dimensions to discover regionalized cellular groups and their co-expression similarity. We identified 3D networks of intact ENS surrounding the gut and predicted cellular connectivity properties based on the stage. Spatial variable genes, such as *hoxb5b*, *hoxa4a*, *etv1*, and *ret*, were regionalized along gut axes, suggesting they may have a precise spatiotemporal control of ENS development. The application of SGA to ENS development provides new insights into its cellular transcriptional networks and interactions, and provides a baseline data set to further advance our understanding of gut neurodevelopmental disorders such as Hirschsprung disease and congenital enteric neuropathies.

## Introduction

The ENS is a complex network of neurons and glial cells embedded within and surrounding the entire length of the gut. It is responsible for coordinating, regulating, and modulating various functions of the digestive system such as peristalsis, gut hormone secretions, and water balance homeostasis ^1,2^. During animal evolution, the ENS is thought to have arisen independently of and prior to the central nervous system (CNS), gaining its designation as the “first brain”. Its ability to function autonomously without direct CNS input further supports this argument ^3^. Moreover, as the primary modulator of gut microbiota ^4^, the ENS plays a crucial role in coordinating the gut-brain axis (GBA), associating microbiota alterations with neurological diseases ^5^.

During development, the vertebrate ENS is derived primarily from a transient unique population of dynamic, multipotent, highly migratory stem cells called neural crest cells (NCCs) ^6^. These NCCs delaminate extensively from the neural tube and migrate to different embryonic locations, where they differentiate into an array of different cell types, including melanocytes, cartilage, bone, smooth muscle, and peripheral nerves ^7^. A subpopulation called vagal NCCs migrate into the foregut, at which this point they are referred to as enteric neural progenitors (ENPs), or sometimes enteric neural crest cells (ENCCs), where there are able to migrate and differentiate into enteric neurons or enteric glia ^8^. Genes encoding proteins such as transcription factors and signaling receptors that are known to be required for controlling ENS developmental processes include *Sox10*, *Foxd3*, *Hoxb5*, *Phox2b*, *Ret*, and *Gfra1,* among many others ^9,10^. In addition, understanding of zebrafish ENS differentiation has made great strides over the past several years, such that we understand aspects of early ENS formation from a gene regulatory perspective, ranging from 3 to 7 days post fertilization (dpf) (Uribe, 2024). However, we still do not fully understand how ENPs differentiate and regionally pattern along the gut. During ENS development, several diseases can arise from genetic mutations, disrupted molecular signaling, or environmental triggers. The most common is Hirschsprung Disease (HSCR), caused by the incomplete migration, proliferation, survival, or differentiation of ENPs along the gut ^11^. Thus, the premise for elucidating how the ENS differentiates along the gut is strong.

Toward that end, the application of ENS single cell RNA-sequencing (scRNA-seq) technology in various cell and animal models has provided valuable insights into neuronal diversity, glial cell populations, and developmental dynamics ^12–14^. Specifically, zebrafish ENS research has benefited greatly from the scRNA-seq approach revealing several major vertebrate discoveries such as the identification of novel cell populations, a binary neurogenic branching of ENS differentiation, and a glial source of enteric progenitors (ENPs) ^15–17^. This was done by leveraging key zebrafish advantages such as high fecundity, genetic conservation with humans, shared molecular pathways, optical transparency, rapid gene editing, and embryonic development ^18^. Particularly, the zebrafish ENS displays a less complex architecture compared to mammals, lacking the submucosal plexus and displaying scattered enteric neurons in contrast with the clustering of ganglia, making it easier to dissect core mechanisms ^19,20^.

Nonetheless, to understand ENS network complexity during development, preserving its intact spatial context at a single cell resolution is required to glean further insights. Recent efforts have been made to process multiplex spatial mRNAs in zebrafish tissues ^21^, and spatiotemporal dynamics of the developing ENS in zebrafish has been done in 2-dimension (2D) guts in coordination with scRNA-seq ^22^. To leverage the spatial and expression dimensions entirely we used spatial genomic analysis (SGA), a cutting-edge method that enables the detailed examination of gene expression patterns in the positional and network context of cells relative to one another and the tissue ^23,24^.

In this study, we developed an SGA pipeline for the developing zebrafish ENS that consisted of combining larval heterogeneity over time along with their 3D spatial and transcriptional data utilizing known markers that encompassed the developmental trajectory of the ENS. By integrating early and late developmental ENS stages, we identified the clustering of cell communities that span specific regions along the gut. The different groups were identified by their differential gene marker expression, and the intercellular networks between different stages displayed changes in node strengths, proximity and density. In addition, we found expression correlations between different genes and the identification of spatial genes based on XYZ position of the gut and the developmental stage.

## Results

### 3D imaging of the developing ENS from the whole gut level using sequential multiplexed whole mount HCR in Slide

To capture the 3D spatial gene expression profiles of differentiating ENS cells along the entire gut length, sequential multiplexed whole mount hybridization chain reaction (HCR) was utilized on 4 and 7 dpf larval fish that were positionally fixed inside chambered slides (**Figure 1A**). We selected 12 ENS marker genes that encompass the developmental transition from ENPs to enteric neurons in zebrafish (**Table 1**). The full sequential multiplexed HCR detection involved four rounds that detected three gene markers each, with the removal of DNA-based HCR probes and hairpins between each round (**Figure 1B and C**). Confocal imaging of the whole gut at 4 dpf following the first round revealed *phox2bb*, *etv1*, and *oprd1b* mRNA expression among differentiating ENS cells across the foregut, midgut, and hindgut (**Figure 1D**). Through machine learning-based cell segmentation and microscopy image analysis (IMARIS), we positionally identified the ENS cells, enabling the extraction of quantitative data, including gene expression and 3D (XYZ) spatial positioning along the gut (**Figure 1E**). In every HCR round, we identified the same ENS cells within the gut, expressing the different ENS markers (**Table 1**) in the gut tissue (**Figure 1F**). From Z-slice views, the ENS cells were observed to distribute circumferentially around the gut tube (**Figure 1G**), as previously shown ^25^. After rounds of HCR, confocal, and imaging analyses, the segmented spatial gene expression data allowed us to visualize the developing ENS along 3D coordinates (**Figure 1H**), and provided metadata for further investigation, as described below.

**Figure 1.**
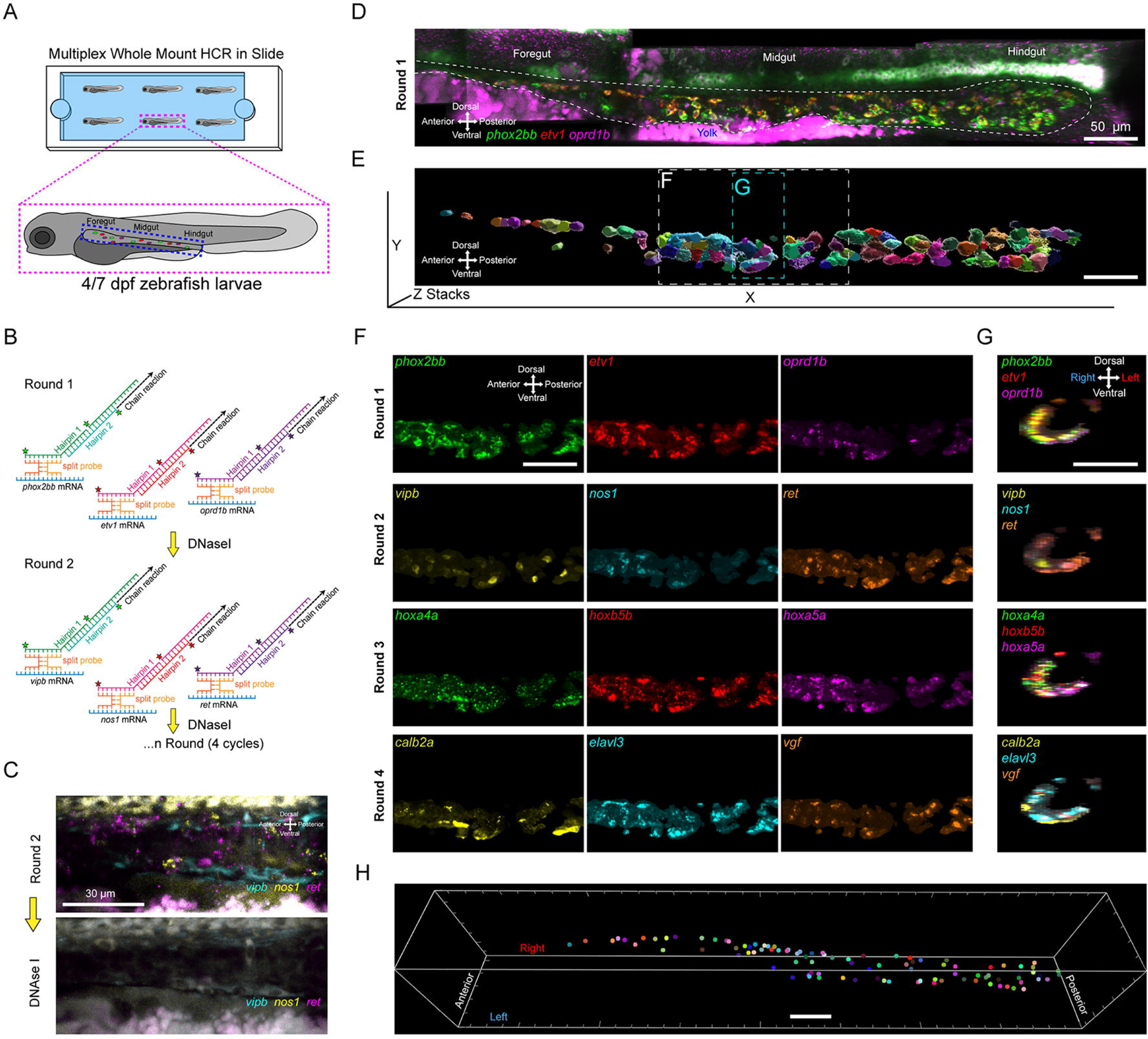
3D imaging of the developing zebrafish gut using Multiplex Whole Mount HCR in Slide. **A)** Zebrafish larvae at 4 and 7 dpf were fixed in a chambered silanized slide to detect transcripts using HCR in intact tissue. **B)** HCR technology is based on DNA split probes that, in the presence of a target mRNA, unwind fluorescently labeled DNA hairpins to induce a chain reaction between them. Each imaging round detected three mRNAs. In between each round, there was a DNase I treatment to degrade the DNA probes and hairpins to start the next round for a total of 4 cycles. **C)** Maximum intensity confocal z-stack projection of the zebrafish ENS before and after the DNAse I treatment, effectively removing the DNA probes and hairpins for *vipb*, *nos,* and *ret.* **D)** Stitched longitudinal maximum intensity confocal z-stack projection of a whole 4 dpf ENS detecting the first round of HCR for *phox2bb*, *etv1* and *oprd1b*, with dashed lines around the ENS and the yolk is located at the ventral region. **E)** Longitudinal view of segmented ENS cells represented in 3D, with each one colored differently. Dashed rectangles denote the longitudinal or transverse planes for Figure 1F and G. Scale bar length is 50 µm. **F)** 4 dpf larvae longitudinal or **G)** transverse plane maximum-z-stacks of the midgut region showing 4 different rounds of HCR with the different fluorescent signals for each of the probes shown in split panels. **H)** 3D map of segmented cells represented as colored points from a dorsal view containing their coordinates and gene expression metadata used for the SGA.

**Table 1.** ENS marker genes used for SGA in this study.

| **Zebrafish gene symbol** | **Zebrafish gene name** | **mRNA Accession used for HCR probe design** | **ENS marker type** | **References** |
| --- | --- | --- | --- | --- |
| *phox2bb* | *paired like homeobox 2Bb* | NM_001014818.1 | ENP, Differentiating enteric neuron, Enteric neuron | 10,15 |
| *etv1* | *ETS variant transcription factor 1* | XM_005157634.4 | Differentiating enteric neuron, Enteric neuron | 37,56 |
| *oprd1b* | *opioid receptor, delta 1b* | NM_131258.4 | Differentiating enteric neuron | 15,38 |
| *vipb* | *vasoactive intestinal peptide b* | NM_001114555.1 | Differentiating enteric neuron | 15,57 |
| *nos1* | *nitric oxide synthase 1* | NM_131660.1 | Differentiating enteric neuron | 15,58 |
| *ret* | *ret proto-oncogene receptor tyrosine kinase* | NM_181662.2 | ENP, Enteric neuron | 10,35 |
| *hoxa4a* | *homeobox A4a* | NM_131535 | ENP | 15,59 |
| *hoxb5b* | *homeobox B5b* | bc078285.1 | ENP | 15,49 |
| *hoxa5a* | *homeobox A5a* | NM_131540.1 | ENP | 15 |
| *calb2a* | *calbindin 2a* | NM_200718.1 | Differentiating enteric neuron | 16 |
| *elavl3* | *ELAV like neuron-specific RNA binding protein 3* | NM_131449 | ENP, Differentiating enteric neuron | 15,20 |
| *vgf* | *VGF nerve growth factor inducible* | XM_003198950.5 | Differentiating enteric neuron | 16,38 |

### Spatial genomic analysis (SGA) integrates larval ENS heterogeneity during development

While it is appreciated that the differentiating zebrafish ENS exhibits regional, temporal, and cellular heterogeneity ^15,16,26^, the 3D spatial details of such cellular heterogeneity and regional gene expression have not been thoroughly explored across stages. In order to capture the range of variability of these important biological mechanisms we integrated several larvae spanning early (4 dpf) and late (7 dpf) ENS differentiation (**Figure 1**) through integrative SGA using the harmony algorithm and the Giotto SGA pipeline (**Figure 2C**) ^27,28^. The expression data of the 12 ENS gene markers and the 3D spatial coordinates of all larval ENS cells from the early and late timepoints were used to generate a low dimensional representation UMAP (Uniform Manifold Approximation and Projection) (**Figure 2A**). These cells were clustered together based on the Leiden community detection algorithm ^29^, creating 12 different ENS groups (**Figure 2B**). Groups 1 to 4 were present in the 4 dpf larvae, being more noticeable in larval fish 1 and 2, and less in larval fish 3 and 4, highlighting the existence of heterogeneity between larvae from the same stage (**Figure 2A, B, D’-G’**). However, cells from groups 1 and 4 were located mostly at the hindgut ENS region (**Suppl Fig 1**). 7 dpf larval ENS was comprised mainly of groups 5 to 12 (**Fig 2B, H, I**) and its location along the gut was mostly dispersed. Moreover, the early and late larval groups exhibited differential gene expression signatures, as noted in the similarity heat plot where groups 1 to 4 were divided from 5 to 12 (**Figure 3A**). When we compared the multidimensional data, with dimensions defined by gene expression and spatial coordinates, we observed that the 4 dpf heterogeneity of larval fish 3 and 4 was in between that of the other early larvae 1 and 2, and the 7 dpf larval fish 1 and 2 (**Figure 3B**). In the UMAP visualization colored by fish stage, it was also clear that the top left quadrant contained the early larval ENS cells, the top right quadrant a combination of both, and the bottom middle part most of the late stage ENS cells (**Figure 3C**).

**Figure 2.**
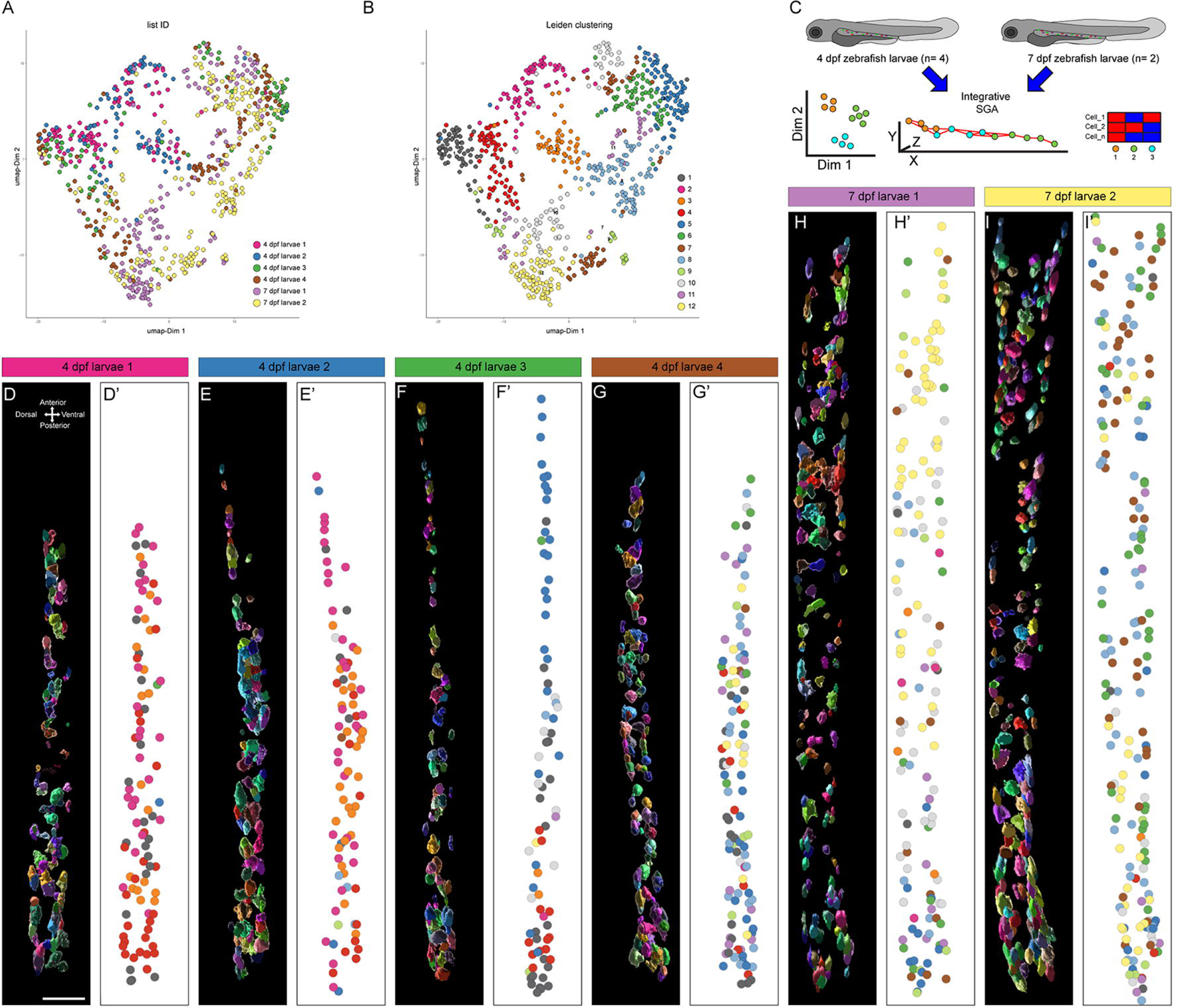
Integrative Spatial Genomic Analysis (SGA) of the early and late developing zebrafish ENS. **A)** UMAP plot integrating different larvae from early 4 dpf and late 7 dpf stages denoting each larva by color. **B)** UMAP plot reveals 12 distinct groups based on the Leiden algorithm. **C)** Diagram of the SGA pipeline that used 2 larval stages for the integration of the spatial and expression data. **D-G)** Spatial plot longitudinal view of segmented ENS cells represented in 3D, with each one colored differently for the 4 dpf and for the **H, I)** 7 dpf stage. **D’-G’)** Spatial plots showing longitudinal 2D visualization of the 4 dpf cells and for the **H’, I’)** 7 dpf stage colored based on the Leiden 12 distinct groups.

**Figure 3.**
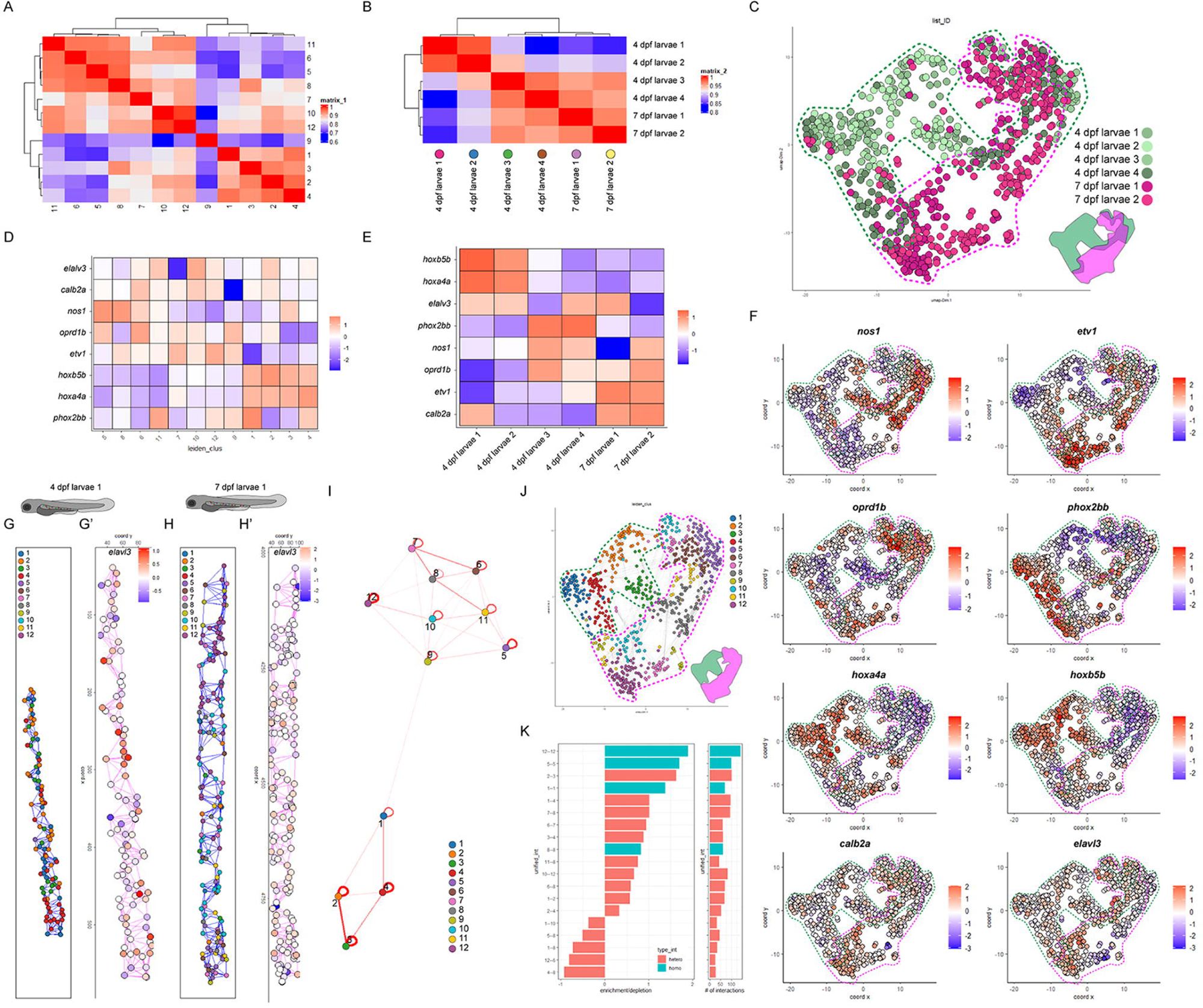
SGA of the developing ENS identified differentially expressed genes and cell networks along the gut. **A)** Heat plot based on gene expression to identify similarities between the different identified groups by the Leiden algorithm. Dendrograms are located at the top and left position representing the hierarchical relationship between groups. **B)** Heat plot based on gene expression data to compare differences between larvae from different stages. Dendrogram to hierarchical cluster larvae are present on the sides. **C)** UMAP plot coloring cells by larvae stage, along with dashed lines in green and pink that surround the regions populated by different cells from different stages. A tiny map on the bottom right side is colored in green and pink. **D)** Heat plot displaying the top DEGs for each Leiden cluster. **E)** Heat plot denoting the top DEGs for the different larvae. **F)** Feature plot expression levels of the DEGs. **G, H)** Spatial plots with colored cells using the cell proximity Leiden network colors from panel I) of a 4 dpf and a 7 dpf, respectively; blue lines denoting the ENS generated network. **G’**, **H’)** Spatial feature plot displaying the expression of the *elavl3* gene in the early and late ENS, respectively, pink lines denoting the ENS generated network. **I)** Cell proximity network representing spatial interactions between different cell types based on their spatial locations within a tissue region; red lines vary in thickness depending on the network strength in connectivity between the different Leiden groups. **J)** UMAP plot with cells colored based on the cell proximity Leiden network. In green and pink are the dashed regions of the early and late larvae and the map. **K)** Bar plot based on the cell proximity scores denoting the enrichment/depletion (left) and a number of interactions (right); pink- and turquoise-colored bars denote the type of heterotypic and homotypic interactions, respectively.

### Differentially expressed genes define clusters during ENS development

The previously observed groups obtained by the Leiden algorithm (**Figure 2B**) were then subjected to analysis for the detection of differentially expressed genes (DEGs) by comparing each cluster against all other clusters with the aim of finding genes that are uniquely or significantly more highly expressed in one cluster compared to the rest. Consequently, we identified the top-ranked marker genes for each cluster (**Figure 3D**). For example, Cluster 4 had a DEG combination based on relatively high expression of *hoxb5b*, *hoxa4a,* and *phox2bb* and low expression of *oprd1b* and *etv1*. Groups 1 to 4 shared high expression of *hoxb5b* and *hoxa4a*; these groups, as previously mentioned, are part of the 4 dpf ENS, which can also be noted in the DEGs comparison by larvae (**Figure 3E**). In contrast, *etv1* was shared in groups 5 to 12, which was expressed mainly in the late larvae (**Figure 3D and E**). All of these DEGs were indeed expressed at different levels on the UMAP expression 2D dimensional reduction plots (**Figure 3F**) and were localized based on the developmental ENS stage in the UMAP plot (**Figure 3C**) but also positionally along the ENS stages, as for example with the *elavl3* gene (**Figure 3G’ and H’**).

### Differentiating ENS network structure during development

To assess and quantify cell-cell interaction enrichment based on the spatial proximity of the different cells, we generated spatial networks of the early and late larvae using the Giotto SGA package, which enables systematic spatial gene association analysis to identify enriched cellular interactions. Overall, there were more interaction network nodes and cells in the 7dpf larvae compared to the 4 dpf, as expected in a more mature ENS (**Figure 3G, H and J**). The cell proximity network showed division between the early and late ENS groups and the strength of nodes between clusters (**Figure 3I**). For example, the cell interactions between groups 2 and 3 were stronger than others (**Figure 3G, I**). In contrast, while the modeled strength in between clusters of the late ENS was not as strong, there were more interactions, resulting in significant interconnectivity (**Figure 3H, I**). This also can be noted in the cell proximity bar plot (**Figure 3K**) for groups 2 and 3 from the 4 dpf larvae that have higher heterotypic interactions, involving interactions between different cell types, as opposed to groups 12 and 5 from the 7 dpf ENS with mostly homotypic interactions, where cells of the same type interact more frequently, but both with enriched networks (**Figure 3I**). Thus, the DEGs and cell networks suggest precise temporal and spatial control of ENS gene expression during ENS development along the gut.

### ENS spatial gene detection during development

Delineating complex spatial patterns of gene expression along the gut is essential for understanding ENS organization, function, and development. To that end, we identified genes that had similar spatial expression patterns along the developing zebrafish larval gut (**Figure 4A**). A heat map for the spatial correlation between marker genes revealed a partition of 5 clusters (**Figure 4A**, colored bar), which showed *nos1* and *vipb* (group 1); *etv1*, *oprd1b* and *calb2a* (group 2); *hoxa4a* and *hoxb5b* (group 3); *phox2bb* (group 4); *ret*, *hoxa5a*, *elavl3*, and *vgf* (group 5) forming a spatial correlation network. Interestingly, *phox2bb* was alone in group 4, which may be explained by the broader relative expression compared to other markers.

**Figure 4.**
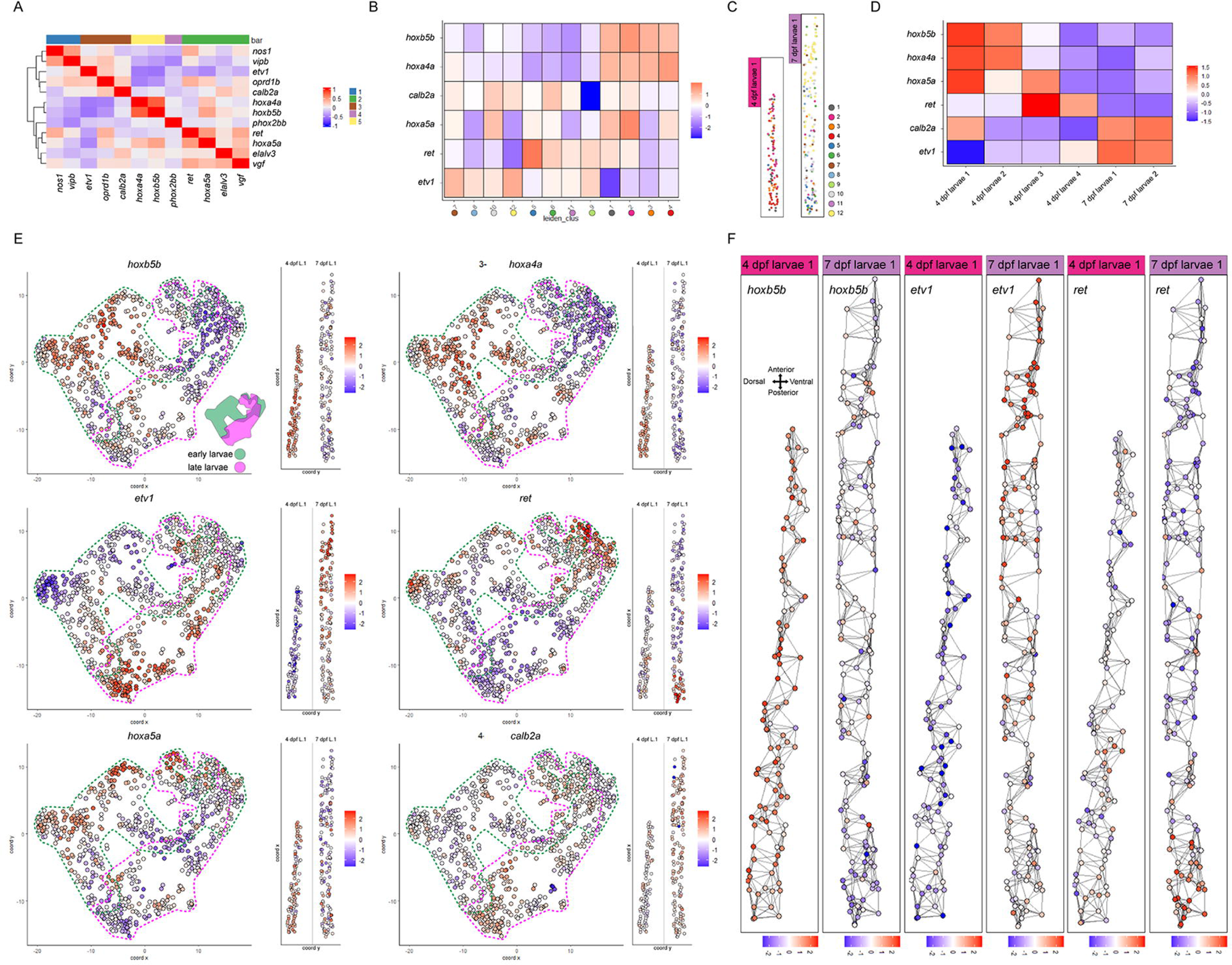
Identification of ENS Spatial Genes during development. **A)** Heatmap showing the correlation of spatial expression patterns among the 12 expressed genes in the dataset. Groups of genes with similar expression profiles were clustered into 5 spatial co-expression modules, which are indicated with different colors on top. **B)** Heat plot identifying the top spatial genes for each cell group from the Leiden algorithm. **C)** Spatial plots coloring cells based on the Leiden clustering along the gut. **D)** Heat plot of the top spatial genes present in the larval ENS from both early and late stages. **E)** UMAP feature and spatial plots color the enriched location of each top spatial gene. Dashed lines in green and pink surround the regions populated by different cells from different stages, and the minimap is shown only in the *hoxb5b* panel. **F)** Spatial network of the 4dpf larval fish 1 and the 7dpf larval fish 1 for the *hoxb5b*, *etv1* and *ret* spatial genes.

Next, we identified the top spatially variable genes (SVGs) along the larval gut. SVGs are genes whose gene expression levels exhibit non-random, informative spatial patterns ^30^. Based on the Leiden clustering, we found that 3 *hox* genes ranked in the top 6 alongside *ret*, *calb2a,* and *etv1* (**Figure 4B**). *hoxb5b* and *hoxa4a,* as mentioned previously were also DEGs and were relatively enriched along the 4 dpf ENS but not in the 7 dpf ENS (**Figure 4B and C**). This was confirmed also in the spatial gene expression heat map of the individual fish larvae from different stages (**Figure 4D**). In agreement, the UMAP dimensional and 2D spatial plots showed the expression of the *hox* genes with more granular detail (**Figure 4E**). Interestingly, *hoxa5a* and *ret* were identified as top spatial genes but didn’t show up as DEGs; this may be because the spatial pattern identification was created on a network-based detection algorithm that captures relationships different from the DEGs analysis. By looking in detail at the UMAP and 2D spatial plots of the top spatial genes we found interesting patterns. For example, the *etv1* gene had increased localization at the late ENS stage and there was a preferential regional enrichment in the foregut and midgut (**Figure 4E and F**). A similar spatial pattern was present with *calb2a* and was expected since they are from the same spatial correlation group 2 (**Figure 4A**). The spatial expression of *hoxb5b* was relatively distributed throughout the 4 dpf ENS but largely became restricted to the foregut and midgut by 7dpf (**Figure 4F**). Another type of spatial arrangement was with the *ret* gene, which had a predominant regional enrichment along the hindgut at early and late stages (**Figure 4E and 4F**).

## Discussion

Our study provides a systems-level spatiotemporal transcriptional profiling of the zebrafish ENS during development, which identified a 3D network of differentiating ENS cells. We used these spatial datasets to identify distinct cellular groups, differentially expressed genes, cellular connectivity, and spatial genes. Recently, different spatiotemporal analyses of human and mouse intestines under different conditions and stages have been able to capture enteric neuron patterns at multi-cellular resolution ^31^, making it possible to recognize neuronal markers such as human PHOX2B, NOS1 and VIP; and mice Uchl1, Pas, and Stmn2; however ENS compartmentalization was not further investigated. While a previous study has explored single-cell transcriptome heterogeneity in the zebrafish ENS spatially ^22^, to our knowledge, ours is the first study that combines 3D (XYZ) spatial and transcriptional data from multiple larval stages to generate cellular clusters and subsequent SGA. This not only enabled the identification of ENS groups but also generated a detailed neuronal network that identified differences in the interconnectivity based on expression, position and stage of the ENS.

Regionalization of the developing ENS has been studied in several models, finding patterns along the entire gut ^32^. Longitudinally, nitrergic neurons (nNOS+/Hu+) have been found proportionally to be slightly higher distally as compared to the proximal colon in adult mice ^31^. Those results are analogous to our study where co-expression of the *elavl3* and *nos1* markers are increased at the hindgut region in early and late stages (**Figure 3D, F**) being part of groups 3 and 5 respectively (**Figure 2B, D’-I’**). Specifically, in adult zebrafish and larvae, neurochemical codes have been established based on combinatorial markers along the gut regionalized in three longitudinal sections: foregut, midgut and hindgut ^33^. In general, our study agrees with their findings at 4 dpf where we measured co-expression of *elavl3* with *nos1* and *calb2a* in our DEGs with a tendency to increase in the later stages (**Figure 3D-F**). In addition, the *nos1* reduction at the foregut was observed in our spatial data for groups 7 and 12 (**Figure 3F, J and H**). Two other specific ENS subtypes that have been reported throughout the 6 dpf larvae are ANB (Adcyap1b/Nos1/Vipb) and ANVB, (Adcyap1b/Nos1/Vip/Vipb) with expression in the foregut and midgut ^22^. Our study detected the existence of a *nos1*/*vipb* spatial correlation (**Figure 4A**) and found that only *nos1* was a DEG (**Figure 3D**) but not an SVG (**Figure 4B**), confirming that this population is broadly located along the gut at 7 dpf.

One of the most studied ENS genes during development is the *ret* gene as it is involved in processes like proliferation, migration and differentiation of ENPs. In our study, the *ret* gene didn’t emerge as a DEG but was one of the top SVGs with a spatial patterning towards the ENS wavefront (posterior cells of hindgut) (**Figure 3D, E and F; 4B, E and F**). Our results confirm findings where migratory wavefronts of ENPs expressing *ret* populated the gut ^34^ and its loss produced fewer ENPs that failed to reach the hindgut ^35^, suggesting that *ret* is important for ENS tissue architecture and spatial organization.

An interesting finding in our study was the contrasting spatial difference of *ret* and *etv1*. During the 7 dpf larvae stage, while *etv1* was a SVG along the foregut and midgut, *ret* was a hindgut SVG (**Figure 4E and F**). In the mouse Pacinian corpuscles, a type of sensory nerve, there is a RET-ETV1-NRG1 pathway important for its development, where Ret is required for the maintenance of Etv1 expression ^36^. In addition, a GDNF-Gfra1-Ret-Etv1 pathway analysis of the ENS has shown that Ret-Etv1 expression is required for the differentiation of enteric nitrergic neurons ^37^. The spatial *ret*-*etv1* binarization found in our study suggests the existence of 3D regional differentiation patterns along the zebrafish ENS.

Previously, our group identified *vgf* and *oprd1b* genes to be required for zebrafish ENS development ^38^. The *vgf* gene that codes for a neuroendocrine neuropeptide ^39^ wasn’t in the top spatial or differentially expressed genes along the gut (**Figure 3D, E; 4B**), suggesting its broad effect along the ENS that may start earlier than the 4 dpf stage, and continue to 7 dpf. This agrees with a *Bap1* Cre-Lox knockout mouse model starting at E8.5, where ENP differentiation was impaired and induced an mRNA increase of *Vgf* at postnatal day P5 ^40^, suggesting an early role in neuronal survival, differentiation, and neurite outgrowth ^41^, which are essential processes in the formation of the complex neural networks within the ENS. As part of the opioid receptor family, the *oprd1* gene is linked to opioid receptors, whose general functions in the adult ENS include regulating digestive motility, transit, secretion, and fluid transport ^42,43^. However, opioid receptors’ functions during ENS development remain mostly a novel research area. Our study identified that as a DEG *oprd1b* was specifically upregulated in the late larval ENS groups 5 and 6, where mature enteric neurons are expected to be along the gut (**Figure 3D, E and J**). The fact that opioid receptor encoding transcripts are spatially present in the developing ENS suggests the presence of an opioid endogenous signaling network that is important in the formation of functional neuronal circuits, as it has been shown with the opioid receptors having a role in brain neuroplasticity ^44^.

In the case of Hox genes, it’s known that they are drivers of cell specification, neuronal migration, cell survival, axon guidance and dendrite morphogenesis during the nervous system development ^45^. Spatiotemporal histochemical expression in the mouse and human developing gut have identified that HuC/D+, SOX10+ cells co-expressed several Hox genes such as Hoxa2, Hoxb4 and Hoxd1. Specifically, Hoxa4, Hoxa5 and Hoxb5 were mainly expressed in differentiated neurons at E11.5 and E15.5 in mice ^46–48^. In our zebrafish SGA, we identified *elavl3* co-expression with *hoxb5b* and *hoxa4a* in the early and late stages corresponding to group 2 and 10 respectively (**Figure 3D**). Furthermore, increasing the levels of *hoxb5b* expression expands zebrafish ENPs, influencing ENS development ^49^. In our SGA analysis, we also identified the correlated genes *hoxb5b* and *hoxa4a* to be DEGs (**Figure 3D**) and part of the top SVGs (**Figure 4B**). *hoxb5b* and *hoxa4a* were preferentially expressed at the early ENS differentiation stage 4 dpf (**Figure 3E**), with a spatial tendency towards the hindgut region (**Figure 4E and F**), and with a high spatial correlation (**Figure 4A**); implying that, in part, the spatial organization of the ENS may be dictated by Hox genes co-expression patterns.

Mapping the 3D organization of the ENS has revealed key insights into the architecture and potential roles of distinct cell populations during development. While the ENS has been studied extensively through various approaches, to our knowledge, its developing 3D transcriptional network has not been characterized before, mainly due to technical limitations. In other cases, other neuronal networks have been successfully reconstructed just recently. For instance, the 3D scaffold of the planarian central nervous system (CNS) was resolved, revealing neuron projections connected to muscle structures ^50^. Additionally, a preprint just revealed the molecular 3D domain structure of the mouse olfactory bulb ^51^. In our study, we have shown the evolution of a 3D neuronal network which matures from a small group of ENS cells interacting and surrounding the developing gut, to an increased group of ENS cells with more interconnectivity that surrounds the maturing developing gut (**Figure 3G-K**).

Gaining insights into spatial neuronal networks is essential for uncovering new mechanisms underlying ENS development. Our study introduced a novel framework for analyzing ENS organization in the complete intact gut, offering a valuable tool for further investigations into its structure and function that may help elucidate mechanisms underlying ENS-related disorders, including Hirschsprung disease and the increasing number of other gut diseases.

## Materials and Methods

### Zebrafish husbandry, and larvae collection

This study was carried out in compliance with the Institutional Animal Care and Use Committee (IACUC) guidelines of Rice University. Larvae used in all experiments were obtained from controlled breeding of adult zebrafish to ensure synchronized development. They were maintained at 28°C in standard E3 embryo medium until 24 hours post-fertilization (hpf), after which they were transferred to a 0.003% 1-phenyl 2-thiourea (PTU)/E3 solution ^52^. The embryos utilized were AB WT. Embryos and larvae were removed from their chorions at the developmental stage specified for each experiment.

### Sequential multiplexed whole mount hybridization chain reaction

Larvae zebrafish from 4 dpf and 7 dpf were fixed with 4% paraformaldehyde and positioned permanently on silanized poly-:L Lysine treated slides fitted with Hybriwell sealing system chambers (Gracebio). The larvae were subjected to 4 rounds of hybridization chain reaction (HCR) as previously described ^53^. HCR probes were ordered from Molecular Instruments. In brief, each round hybridized the targeted mRNAs with its HCR probes (Table 1) at 37°C overnight. Subsequently, the probe was washed, and then HCR amplification was done using each gene-specific HCR amplifier with the 488, 546, and 647 fluorophores. In between each round, DNase I (Invitrogen) treatments were done at RT.

### High-content semi-automated confocal microscopy

Confocal imaging of the ENS was performed using an Olympus FV3000 confocal and FluoView software (2.4.1.198), with an Olympus 20.0X objective (UPLXAPO2X Oil). Multi-Area-Time-Lapse (MATL) option was used to create reference maps for each larva per round. Z-sections were obtained to capture the whole thickness of the gut, and acquisition areas were stitched via the Fiji image stitching plugin ^54^ or CellSens (Evident).

### 3D cell segmentation and image curation

Stitched whole gut images were imported into IMARIS 10.2 (Oxford Instruments) and positioned in frame to the first HCR round XYZ coordinates. Afterward, the IMARIS AI-powered segmentation tool was used to identify each ENS cell, and manual curation was used to cut and split surfaces. Cell positions and channel intensity statistics were exported in batch and converted into readable expression and position metadata files for the subsequent SGA analysis.

### Data processing, sample integration, and clustering

SGA was done using the agnostic framework and toolbox Giotto 4.2.1. ^27^ which integrates both R and Python for data processing and analysis using the integrated development environment (IDE) Rstudio (posit). In brief, expression and position metadata was used for each larva of 4 dpf and 7 dpf to create joined objects using “joinGiottoObjects”, then filtration “filterGiotto”, normalization “normalizeGiotto” and principal component analysis “runPCA” was done. To simultaneously account for multiple experimental and biological factors we used the harmony algorithm “runGiottoHarmony”. Afterwards, a nearest-neighbor network based on spatial coordinates of cells was done under the “createNearestNetwork” to perform clustering using “doLeidenCluster”.

### Differential expression, spatial network, and spatial gene detection

For the identification of genes that were selectively expressed in a specific cluster, we used the “findMarkers_one_vs_all” using the scran method ^55^. Spatial and feature-based network construction was done using “createSpatialNetwork” based on the k-Nearest Neighbors Network (kNN) using a value of 6 nearest neighbors based on physical distance. For the spatial detection, we used “Binspect” using the rank method with a minimum number of 5 cell-cell interactions for a hub-cell.

---

**Supplementary Figure 1. Spatial cell diversity of the developing ENS. A)** Spatial integrative plot of the different larvae from the 4 dpf and 7dpf, differently colored. Each larval ENS depicted has its own orientation, shown by axes key. **B)** ENS of the different larvae, colored based on the Leiden clusters (groups). **C)** Spatial plot of the ENS separating each group in each larva based on the Leiden clustering colors. The distance and position of all the cells are in scale to the X coordinates.

## Acknowledgments

We gratefully acknowledge the Rice University Shared Equipment Authority and Dr. Alloysius Budi Utama for providing training, assistance, and access to the IMARIS workstation used for image analysis in this study. We acknowledge Laura Kerosuo and Ceren Pajanoja for their expertise and technical support in the bioinformatics processing and methods standardization of the SGA pipeline. Lastly, we acknowledge Aubrey G. A. Howard IV for his insight, advice, and assistance.

