## Supplementary figures and images for "Whole-gut spatial genomic analysis reveals molecular regionalization of the differentiating zebrafish enteric nervous system"

### Supplementary Figure 1

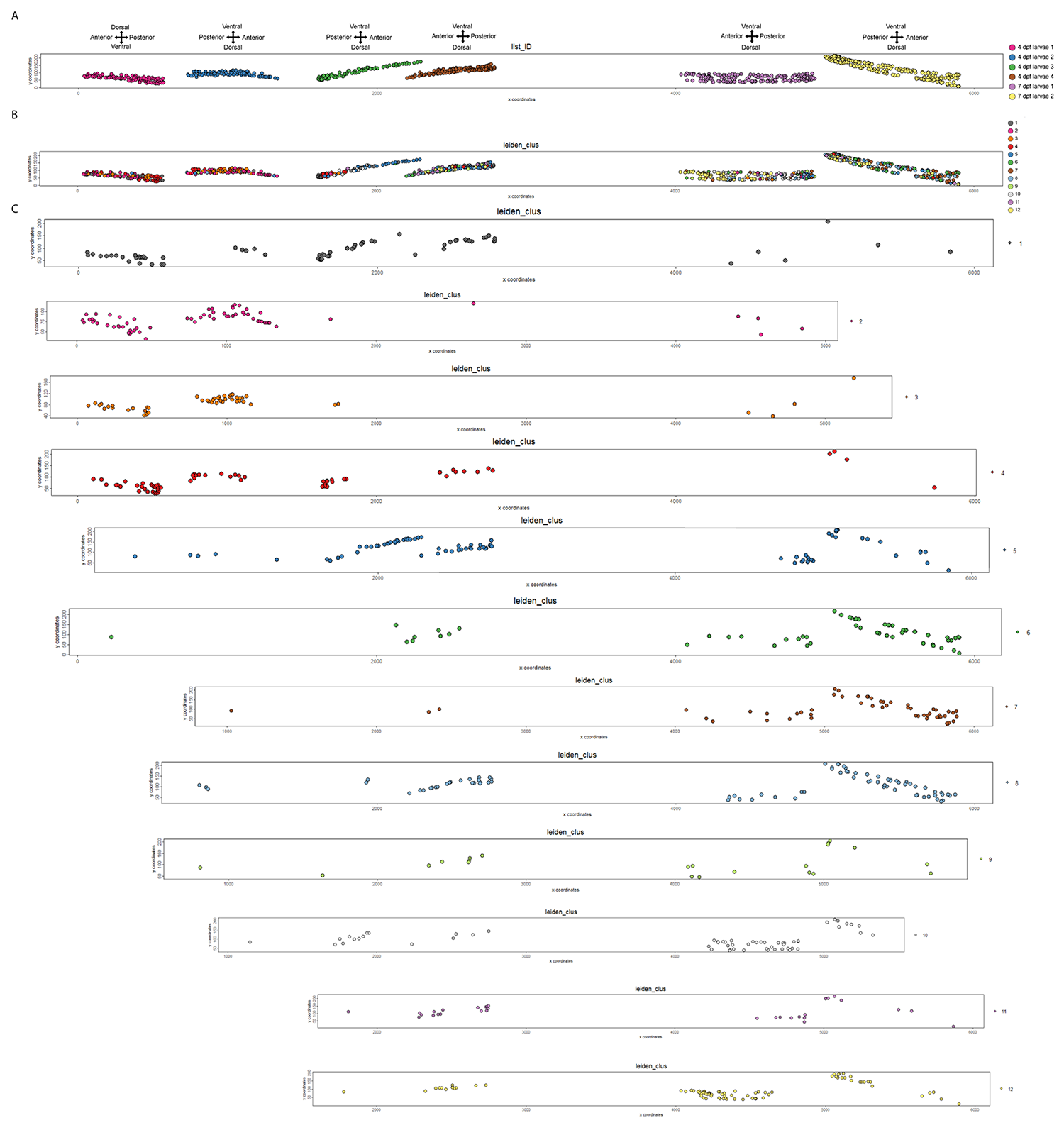
